# A potent SARS-CoV-2 neutralizing antibody recognizing a conserved epitope with broad mutant variant and SARS-CoV activity

**DOI:** 10.1101/2022.02.06.479332

**Authors:** Adam J. Pelzek, Sanam Ebtehaj, James Lulo, Lucy Zhang, Olivia Balduf, Lindsay Dolan, Chaohua Zhang, Shengqin Wan, Gang An, Awo Kankam, Eugene Chan, Shaun P. Murphy

## Abstract

COVID-19 is the deadliest respiratory virus pandemic since 1918 and the latest of several coronavirus epidemics and pandemics in recent years. Despite the unprecedented response by both the government and private sectors to develop vaccines and therapies, the evolution of SARS-CoV-2 variants resistant to these interventions reveals a crucial need for therapeutics that maintain their efficacy against current and future mutant variants. Here we describe a SARS-CoV-2 neutralizing antibody, ABP-310, with potent activity against all variants tested including the Omicron variant. ABP-310 also displays potent neutralizing activity against SARS-CoV, highlighting the conserved nature of the ABP-310 epitope. By targeting a conserved epitope, we believe that ABP-310 has therapeutic promise not only against the current SARS-CoV-2 variants but would be expected to maintain efficacy against future variants and possibly even novel coronaviruses.

---

The coronavirus disease 2019 (COVID-19) pandemic is the deadliest caused by a respiratory virus since the 1918 Influenza pandemic, with over 5.7 million deaths as of February 2022 (Johns Hopkins University COVID-19 Dashboard). The first known case of COVID-19 occurred in Wuhan, China in December of 2019 (Zhou, 2020), the causative agent of which was determined to be a novel human coronavirus subsequently named severe acute respiratory syndrome coronavirus 2 (SARS-CoV-2). SARS-CoV-2 is an enveloped, positive-strand RNA virus in the betacoronavirus genus (Xu, 2020), which also includes the human coronaviruses MERS-CoV and SARS-CoV. SARS-CoV is approximately 80% genetically identical to SARS-CoV-2 and shares angiotensin-converting enzyme 2 (ACE2) as the host receptor to which the spike (S) glycoprotein of both viruses bind (Zhou, 2020). The spike protein exists as a homotrimer on the envelope of the virus with each spike protein monomer comprised of an S1 domain involved in binding to the ACE2 host receptor and an S2 domain, which mediates the fusion of the viral and cellular membranes (Yuan, 2020). The S1 domain consists of an N-terminal domain (NTD), two C-terminal subdomains (SD1 and SD2), and a receptor-binding domain (RBD) in between (Yuan, 2020). The RBD itself is defined as the region between two cysteine residues (C336 and C525) forming a disulfide bridge and dynamically changes confirmation from an “up” state to a “down” state, with only the “up” state being able to bind to ACE2 (Yuan, 2020; Niu, 2021).

While there has been an intense global effort to develop vaccines against SARS-CoV-2 and three vaccines that have received either marketing approval or emergency use authorization (EUA) by the US FDA, from Moderna (Baden, 2021), Pfizer/BioNTech (Polack, 2020), and Johnson & Johnson (Sadoff, 2021) have been shown to be approximately 95% effective for the Moderna and Pfizer/BioNTech vaccines and 67% for the Johnson & Johnson vaccine in preventing infection, their ability to establish lasting long-term protective immunity has not been demonstrated and their efficacy against the Omicron variant is likely weakened (Callaway, 2021; Planas, 2021). In addition, vaccination may not be safe or effective in certain patient populations such as those with vaccine allergies or are immunocompromised. As such, it is critical to develop drugs to prevent and treat COVID-19. Among the most promising class of SARS-CoV-2-targeting therapeutics are neutralizing antibodies (nAbs), many of which can block the spike protein-ACE2 interaction and prevent the virus from infecting host cells. To date, several antibody-based SARS-CoV-2-targeting therapeutics have been given EUA by the US FDA, including COVID-19 convalescent plasma as well as monoclonal nAbs or nAb cocktails from Regeneron, Lilly, and GSK/Vir (FDA emergency use authorizations for COVID-19).

Although nAbs show great promise in the treatment and prophylaxis of COVID-19 (Hurt, 2021), a major potential liability of all nAbs is the evolution of mutant variants resistant to nAb therapy (i.e., escape mutants). Although affecting therapeutic nAbs, the evolution of escape mutants is thought to be largely driven by natural evolution of the virus under selective pressure of the host immune response and not by the selective pressure of antiviral therapies, as acute respiratory infections are normally cleared in immunocompetent individuals receiving such therapies (Holmes, 2021). As a therapeutic involving multiple neutralizing antibodies binding distinct epitopes (i.e., a “cocktail”) would be more resistant to mutant escape as multiple mutation events at different epitopes would typically be required to render a multiple antibody therapeutic ineffective, a cocktail strategy is currently being pursued by multiple biotechnology companies, including a two nAb cocktail approach by Regeneron, Lilly, and Brii. Brii’s cocktail is currently in clinical trials (clinicaltrials.gov; NCT04787211). Regeneron’s cocktail of casirivimab (REGN10933) and imdevimab (REGN10987), currently under EUA, has been demonstrated to reduce the generation of escape mutants *in vitro* compared to either single nAb (Baum, 2020) and this cocktail has been successful in clinical trials (Weinrich, 2021). However, the Regeneron cocktail displays a 20-fold reduction in neutralization potency against the Beta variant of SARS-CoV-2 (Wang, 2021) and a recent study that examined the effect of all possible amino acid mutations in the RBD showed that a single E406W mutation almost completely eliminated the neutralization potency of the Regeneron nAb cocktail (Starr, 2021). Lilly has developed two antibodies used in combination therapy which have received EUA, bamlanivimab and etesevimab. However, one of the antibodies, bamlanivimab, is ineffective against multiple mutant variants including Beta (Wang, 2021) and consequently, its monotherapy EUA was revoked by the US FDA (FDA news release, April 16, 2021). Moreover, the Regeneron and Lilly nAbs, among others, are ineffective against the Omicron variant (Cao, 2021; Planas, 2021), leading the US FDA to limit their use (FDA news release, January 24, 2022). Taken together, these findings suggest that the existing therapeutic neutralizing antibody cocktail strategy alone may not provide adequate resistance to the evolution of escape mutants.

Another promising strategy to combat the emergence of escape mutants is the use of nAbs that recognize conserved epitopes of SARS-CoV-2 shared by other coronaviruses. Since conserved epitopes are believed to occur in protein regions that are essential for some aspect of the viral reproductive cycle, mutations in conserved epitopes are likely to reduce the evolutionary fitness of a virus and would be less likely to become prevalent in the overall viral population (Ekiert, 2009; Ekiert 2011). This conserved epitope approach is being pursued by, among others, Adagio with ADG20 (Rappazzo, 2021) and GSK/Vir with the nAb sotrovimab (Pinto, 2020). The current COVID-19 pandemic is the third public health crisis caused by a novel coronavirus within the last 20 years which suggests the likelihood that other novel coronavirus pandemics may arise in the near future (DeFrancesco, 2020). Neutralizing antibodies with conserved epitope binding and broad activity against not only SARS-CoV-2 but other viruses in the betacoronavirus genus could play a crucial role in the early response to future coronavirus pandemics. Indeed, the US National Institutes of Health has recommended that stockpiling broadly acting nAbs should be a pillar of future pandemic preparedness (Therapeutic Neutralizing Monoclonal Antibodies: Report of a Summit sponsored by Operation Warp Speed and the National Institutes of Health).

The past and potential future vulnerabilities of current nAbs being used under EUA highlights the need for more effective nAb-based therapies. Here we describe the development of a nAb therapeutic approach that uses an antibody, ABP-310, derived from the B cells of a convalescent COVID-19 patient, that binds a conserved epitope and neutralizes the related coronavirus SARS-CoV. We demonstrate that ABP-310 has broad neutralizing activity against recently circulating mutant variants of SARS-CoV-2 including the Omicron variant. In addition, given the conserved epitope binding and neutralization of a related betacoronavirus (SARS-CoV), it is possible that ABP-310 may neutralize future variants of SARS-CoV-2 as well as novel betacoronaviruses yet to evolve.

## Results

### ABP-310 can bind mutant SARS-CoV-2 and SARS-CoV spike protein

Given the failure of many current nAbs to effectively neutralize SARS-CoV-2 mutant variants, we assessed the ability of ABP-310 to bind SARS-CoV-2 spike protein domains bearing a variety of mutations found in variants of concern or those associated with reduced nAb efficacy or enhanced infectivity, such as N234Q, K417N, N440K, K444R, L452R, E484K, F490L, S494P, and D614G (Li, 2020; Grabowski, 2021; Harvey, 2021; Rani, 2021). In addition, the ability of ABP-310 to bind the S1 domain of the spike proteins from MERS-CoV and SARS-CoV was assessed. The expressed sequences of casirivimab, imdevimab, and sotrovimab were used as comparator controls. All antibodies assessed displayed strong binding activity against all SARS-CoV-2 mutants tested, with the exception of a marked reduction in binding of casirivimab to Beta (**Fig. 1**), which is consistent with the previously reported reduced neutralization activity of casirivimab to this variant (Wang, 2021). The only antibodies assayed that displayed binding to either SARS-CoV or MERS-CoV were sotrovimab, a nAb derived from a memory B cell screen of a patient infected with SARS-CoV during the 2003 epidemic (Traggiai, 2004; Rockx, 2008; Rockx, 2010), and ABP-310, both showing binding to SARS-CoV but not MERS-CoV (**Fig. 1**).

**Figure 1.**
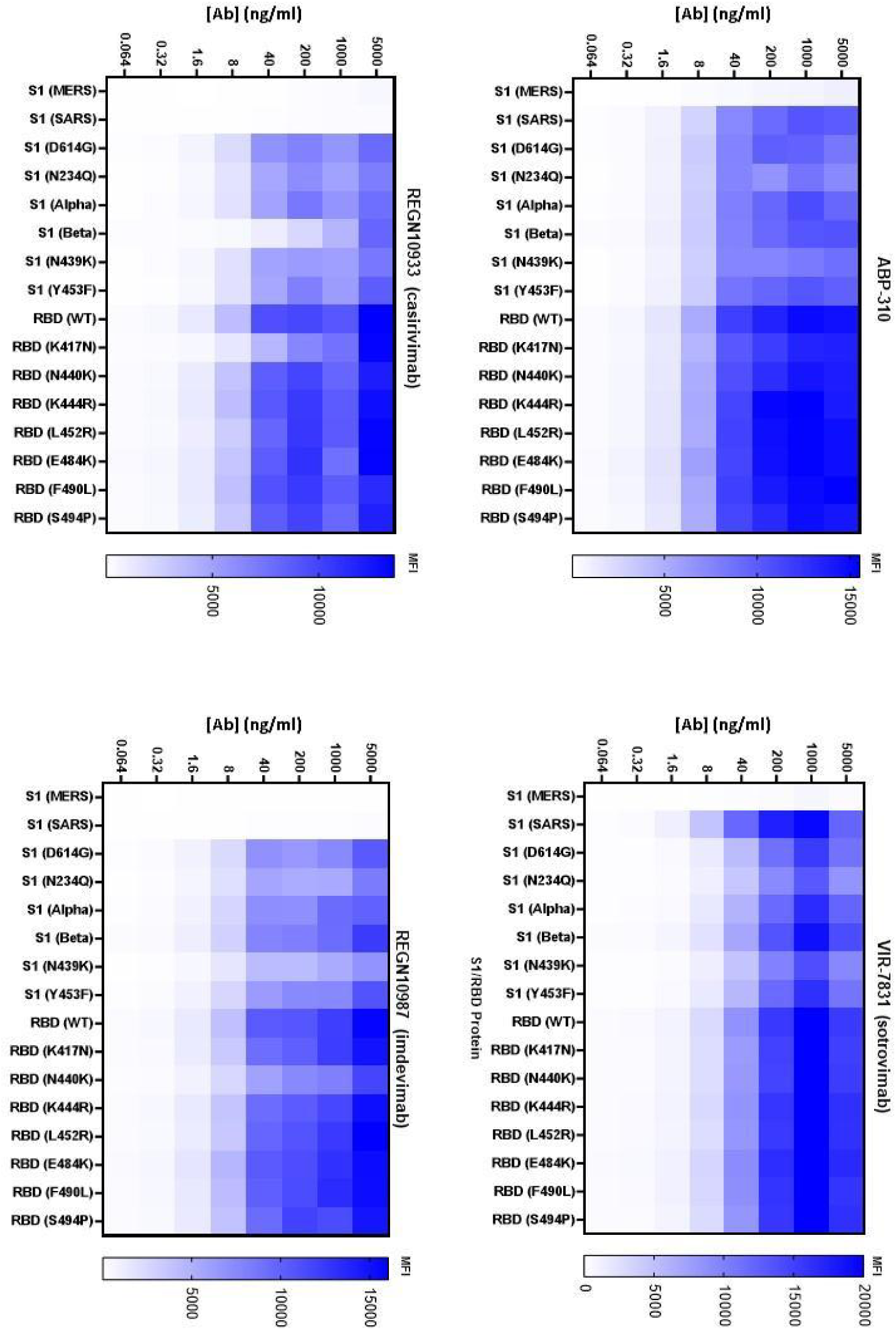
Luminex spike mutant binding. Binding strength as demonstrated by Luminex binding assay. The left vertical axis of each heat map represents the concentration of antibody dilution, the horizontal axis describes the spike protein (with mutations indicated) that are used for each column, and the right vertical axis represents the intensity of binding in median fluorescence intensity (MFI). Data are representative of one independent experiment out of two. All data points were performed in duplicate.

### Characterization of the ability of ABP-310 to block the ACE2-spike protein interaction

As ABP-310 can bind the spike protein RBD (**Fig. 1**), and a major therapeutic mechanism of SARS-CoV-2 nAbs is preventing spike protein binding to ACE2, the ability of ABP-310 to block the spike protein RBD-ACE2 interaction was compared to casirivimab, imdevimab, and sotrovimab. The ability of ABP-310 to block the ACE-2-RBD interaction was demonstrated using an in vitro ACE2 competition assay using the indicated mutant variant spike protein RBDs (**Fig. 2**). The cocktail of casirivimab and imdevimab (REGN-COV2) also blocked the ACE2-spike protein interaction (**Fig. 2**) consistent with published results (Hansen, 2020).

**Figure 2.**
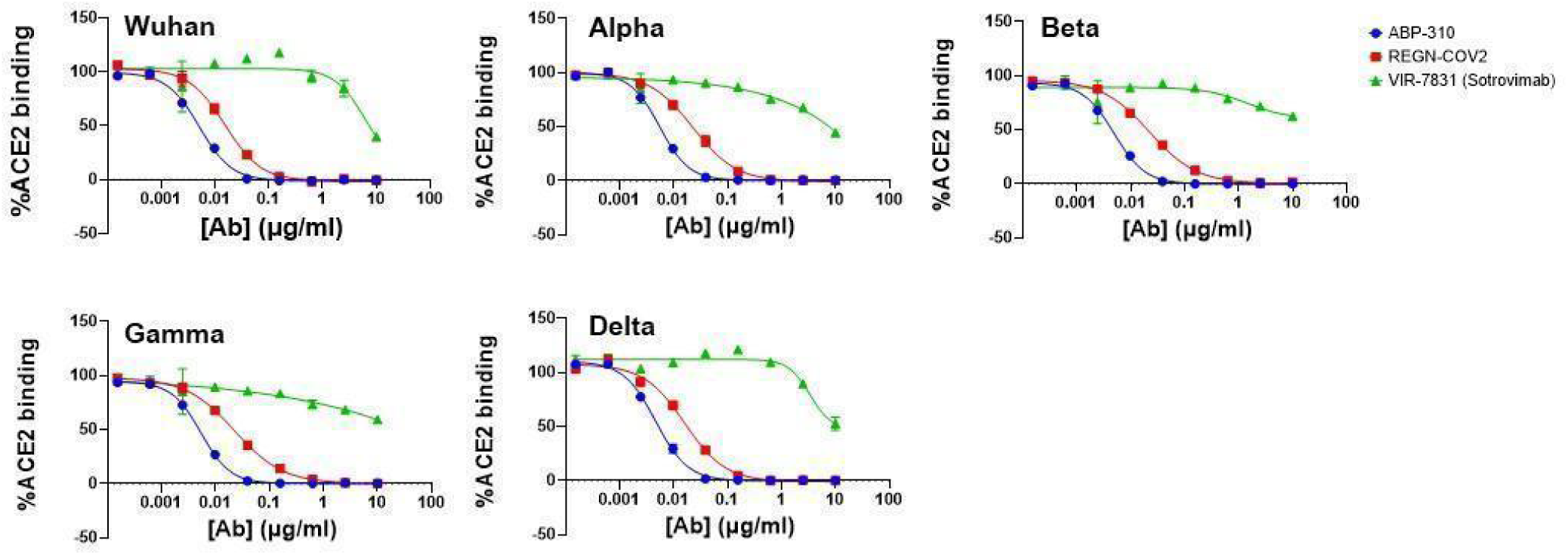
Antibody ACE2 competition. ACE2-competition ELISA using indicated antibodies at concentrations given by the horizontal axis. For REGN-COV-2, casirivimab and imdevimab were added at an equimolar ratio summing to the concentration given on the horizontal axis. A spike protein RBD containing the mutations of the indicated SARS-CoV-2 variants was used. The data for Wuhan is representative of two independent experiments and the data for Alpha, Beta, Gamma, and Delta is from one experiment. All data points are means of duplicates with error bars indicating standard deviation.

Unlike the other nAbs evaluated in this study, sotrovimab did not compete with ACE2 for spike protein binding, consistent with previous studies evaluating S309, the parental antibody to sotrovimab (**Fig. 2**) (Pinto, 2020).

### ABP-310 can neutralize a wide range of SARS-CoV-2 variants and SARS-CoV

All antibodies were assayed for neutralization activity against lentivirus pseudotyped with a variety of SARS-CoV-2 spike proteins including the original Wuhan isolate, all of the current (as of December 2021) WHO variants of concern (Alpha, Beta Gamma, Delta, and Omicron) and major variants of interest as of September 2021 (Epsilon, Eta, Iota) (**Fig. 3, Extended Table 1, Extended Table 2**). Casirivimab displayed a 71-fold reduction in neutralization potency (IC_50_) against the Beta variant (**Extended Table 1**) compared to the original Wuhan virus, consistent with previous data (Wang, 2021), as well as an 89-fold reduced neutralization potency against the Gamma variant (**Extended Table 1**). ABP-310, like sotrovimab, displayed neutralization ability against all SARS-CoV-2 variants tested (**Fig. 3**., **Extended Table 1, Extended Table 2**) with ABP-310 displaying a mean IC_50_ (93 ng/ml) and a mean IC_90_ (407 ng/ml) across variants that were commonly tested and neutralized by all antibodies (Wuhan, Alpha, Beta, Gamma, and Delta) 10.3-fold and 66-fold, respectively more potent than sotrovimab (mean IC_50_: 955 ng/ml; mean IC_90_: 26,812 ng/ml), and 8.5-fold and 20.5-fold, respectively more potent than casirivimab (mean IC_50_: 798 ng/ml; mean IC_90_: 8,348 ng/ml). While ABP-310 exhibited IC_50_s and IC_90_s that are 3.6-fold and 1.6-fold less potent than imdevimab (mean IC_50_: 26 ng/ml mean IC_90_: 201 ng/ml), respectively across variants that were commonly tested and neutralized by all antibodies, neither casirivimab nor imdevimab are able to neutralize Omicron. However, both ABP-310 and sotrovimab neutralize Omicron, with ABP-310 being 2.5-fold more potent than sotrovimab (IC_50_: 0.183 vs 0.453 µg/ml, respectively) (**Fig. 3B, Extended Table 1**). In addition, ABP-310, like sotrovimab, but unlike casirivimab and imdevimab, exhibits neutralization activity against SARS-CoV (**Fig. 4, Extended Table 1, Extended Table 2**), with ABP-310 displaying a 2-fold more potent neutralization IC_50_ for SARS-CoV compared to sotrovimab. As sotrovimab achieved a maximum SARS-CoV neutralization of approximately 75%, a comparison with the IC_90_ of ABP-310 could not be made. It should be noted that while casirivimab displayed detectable neutralization against SARS-CoV (**Fig. 4, Extended Table 1**), it was at least 500-fold weaker in potency compared to ABP-310 (ABP-310 IC_50_: 31 ng/ml *vs*. casirivimab IC_50_: >16,800 ng/ml) (**Fig. 4, Extended Table 1**).

**Figure 3.**
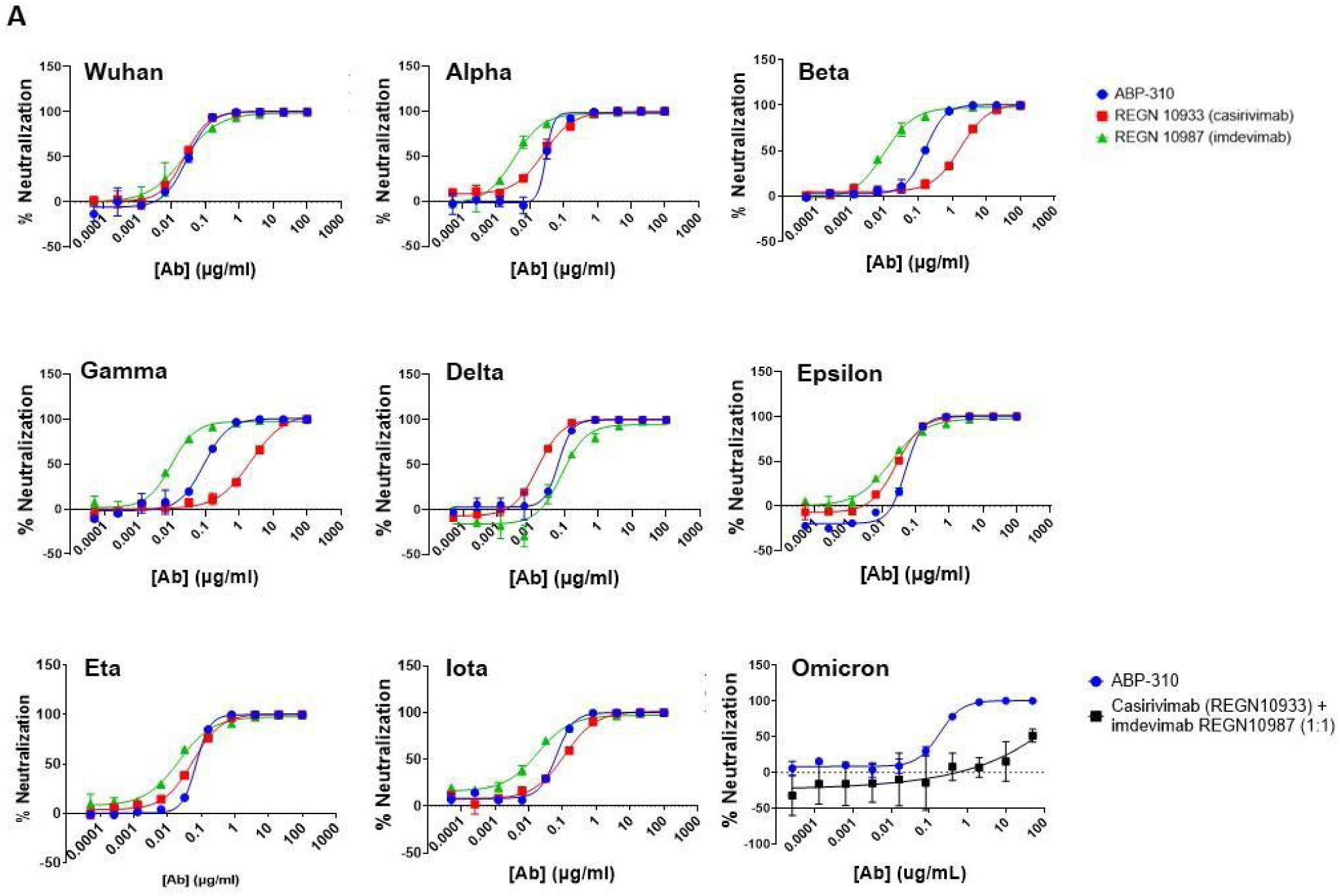
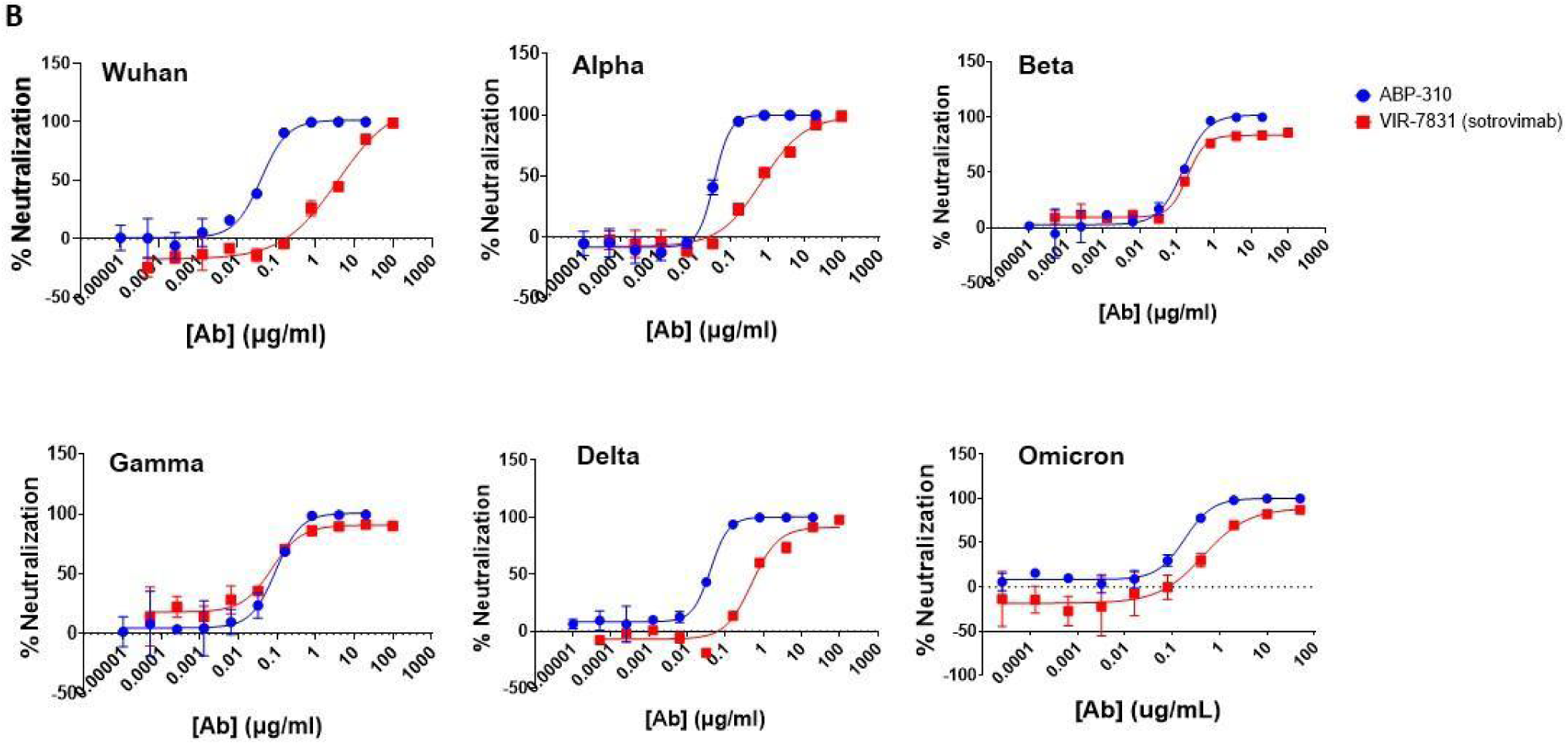
Comparison of pseudovirus neutralization activity of ABP-310 with casirivimab, imdevimab and sotrovimab. Using a SARS-CoV-2 spike pseudotyped viral neutralization assay, the neutralizing activity of each antibody against the indicated SARS-CoV-2 viral variants was determined for ABP-310 and casirivimab and imdevimab (**A**) and sotrovimab (**B**). For the Omicron neutralization assay in (**A**), casirivimab and imdevimab were assayed at an equimolar ratio summing to the concentration given on the horizontal axis. The data are representative of a minimum of two independent experiments. All data points are means of duplicates with error bars indicating standard deviation.

**Figure 4.**
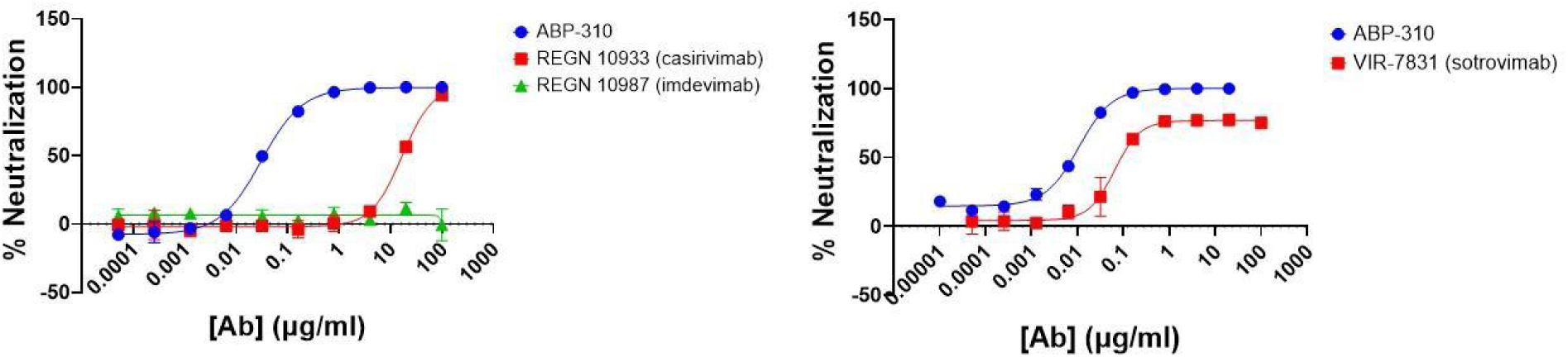
Ability of ABP-310 to neutralize SARS-CoV. Using a SARS-CoV spike pseudotyped viral neutralization assay, the neutralizing activity of each antibody against SARS-CoV was determined. The data are representative of a minimum of two independent experiments. All data points are means of duplicates with error bars indicating standard deviation.

### Conserved nature of the ABP-310 epitope

Given the predicted advantages of a nAb binding a conserved epitope in resistance to mutant escape and efficacy against related viruses, existing or yet to evolve, we assessed the conservation of the ABP-310 epitope by multiple sequence alignment with Lineage B betacoronaviruses (**Fig. 5**). Representative unique spike protein sequences from Lineage B betacoronaviruses (Letko, 2020) were used to generate multiple alignments. SARS-CoV-2 and related variants are identified as clade 1/2 (Letko, 2020), and require ACE2 for cellular entry.

**Figure 5.**
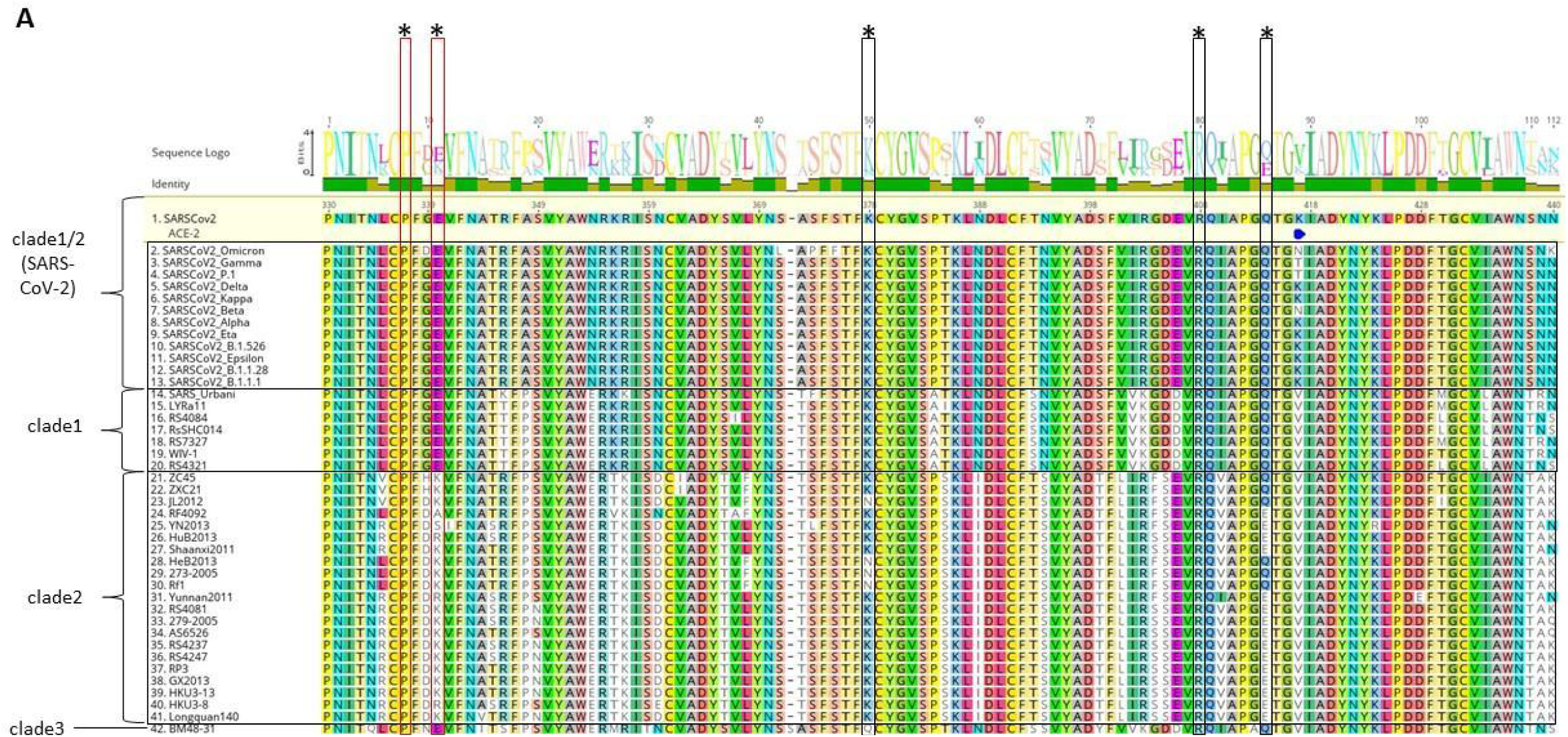
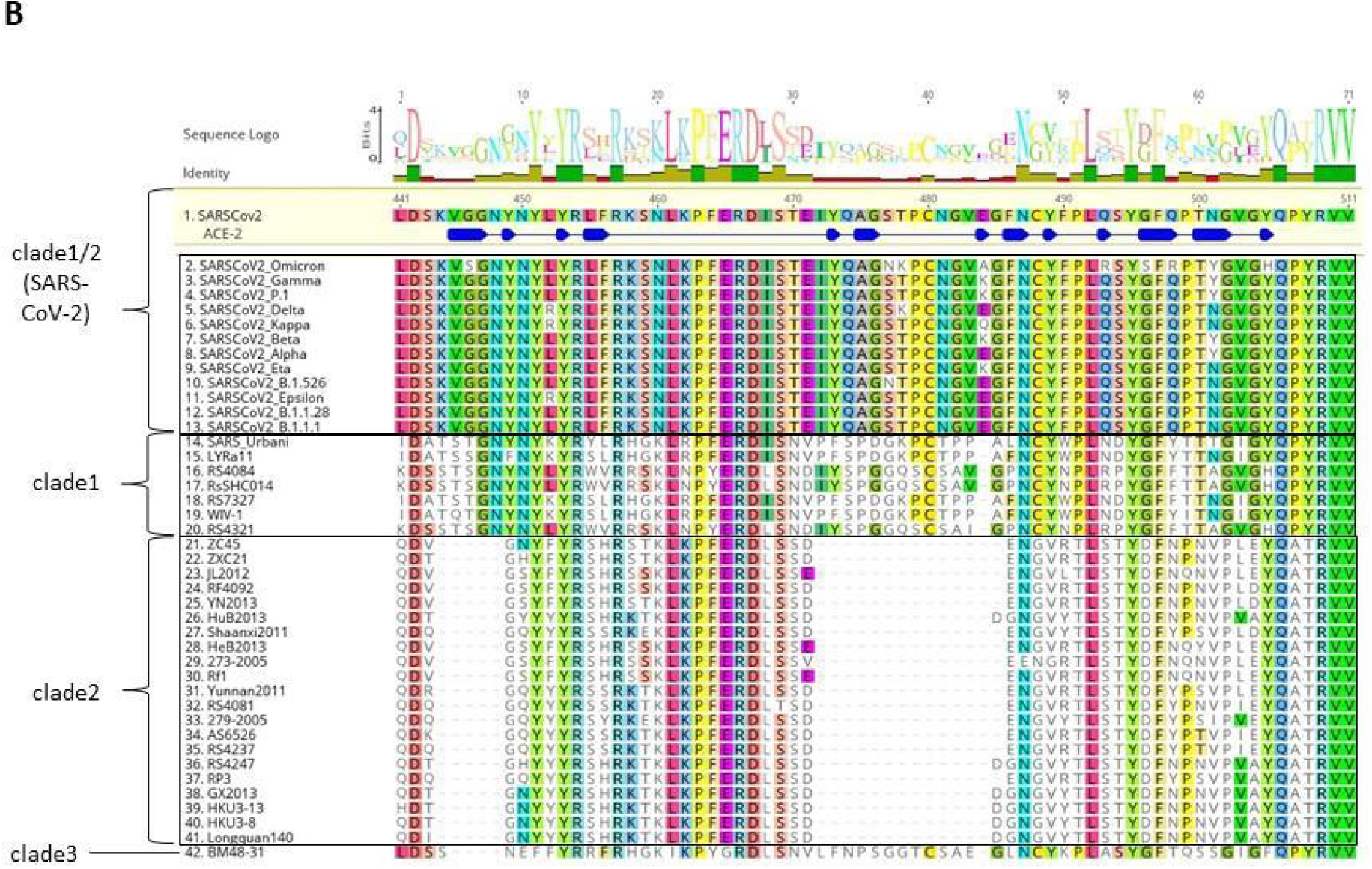
The epitope of ABP-310 is highly conserved. A multiple alignment was made using Clustal Omega in Geneious software with SARS-CoV-2 spike protein (Wuhan isolate; accession YP_009724390.1) used as the reference sequence for designating the position of amino acid residues. Blue bars indicate the binding footprint of human ACE-2. **A**. Amino acids 330-440 of the SARS-CoV-2 spike protein reference sequence and multiple alignment is shown for Lineage B betacoronavirus receptor binding domain (RBD) clades, as previously defined (Letko, 2020), with isolate names shown at left. The logo representation allows qualitative assessment of the frequency of amino acid changes at a given position. Residues labeled as critical for ABP310 or sotrovimab binding by alanine scanning are labeled with an asterisk (*) over black or red boxes, respectively. **B**. Amino acids 441-511 of the SARS-CoV-2 spike protein reference sequence and multiple alignment, with the same requirements as in (**A**). The size and number of the letters in the “Sequence Logo” row in both panels represent the relative proportions of that amino acid at that residue position in the sequences below.

The critical interacting residues for ABP-310 spike protein RBD binding (K378, R408, Q414) were determined by alanine scanning, and those of sotrovimab (P337 and E340) were previously published (Starr, 2021b). Critical interacting residues for both ABP-310 and sotrovimab showed conservation across all SARS-CoV-2 variants of interest, including Delta and Omicron, as well as past variants of concern (**Fig. 5**). In addition, these critical interacting residues are identical in clade 1 RBD isolates as previously defined (Letko, 2020), including SARS, WIV-1, and SHC014. Unlike clade 1/2 and clade 1 viruses, clade 2 virus spike protein RBD sequences do not appear to utilize ACE2 for cellular entry.

The Omicron variant was rapidly elevated to variant of concern status based on the large number of mutations (Stanford University Coronavirus antiviral & Resistance database) that may affect transmissibility, disease severity, or escape from currently available therapeutics (WHO Update on Omicron). No critical binding residues for ABP-310 are mutated in the Omicron variant (**Fig. 6**) and as would be expected, ABP-310 retains neutralizing activity against Omicron (**Fig. 3, Extended Table 1, Extended Table 2**). Several Omicron mutations also coincide with residues that are critical for binding of both of the antibodies in Regeneron’s cocktail, including K417, E484, and Q493 for casirivimab, and N440 and Q446 for imdevimab (Starr, 2021a), (**Fig. 6**), which is consistent with the lack of activity of the Regeneron cocktail against Omicron (**Fig. 3A, Extended Table 1, Extended Table 2**), (Cao, 2021; Planas, 2021).

**Figure 6.**
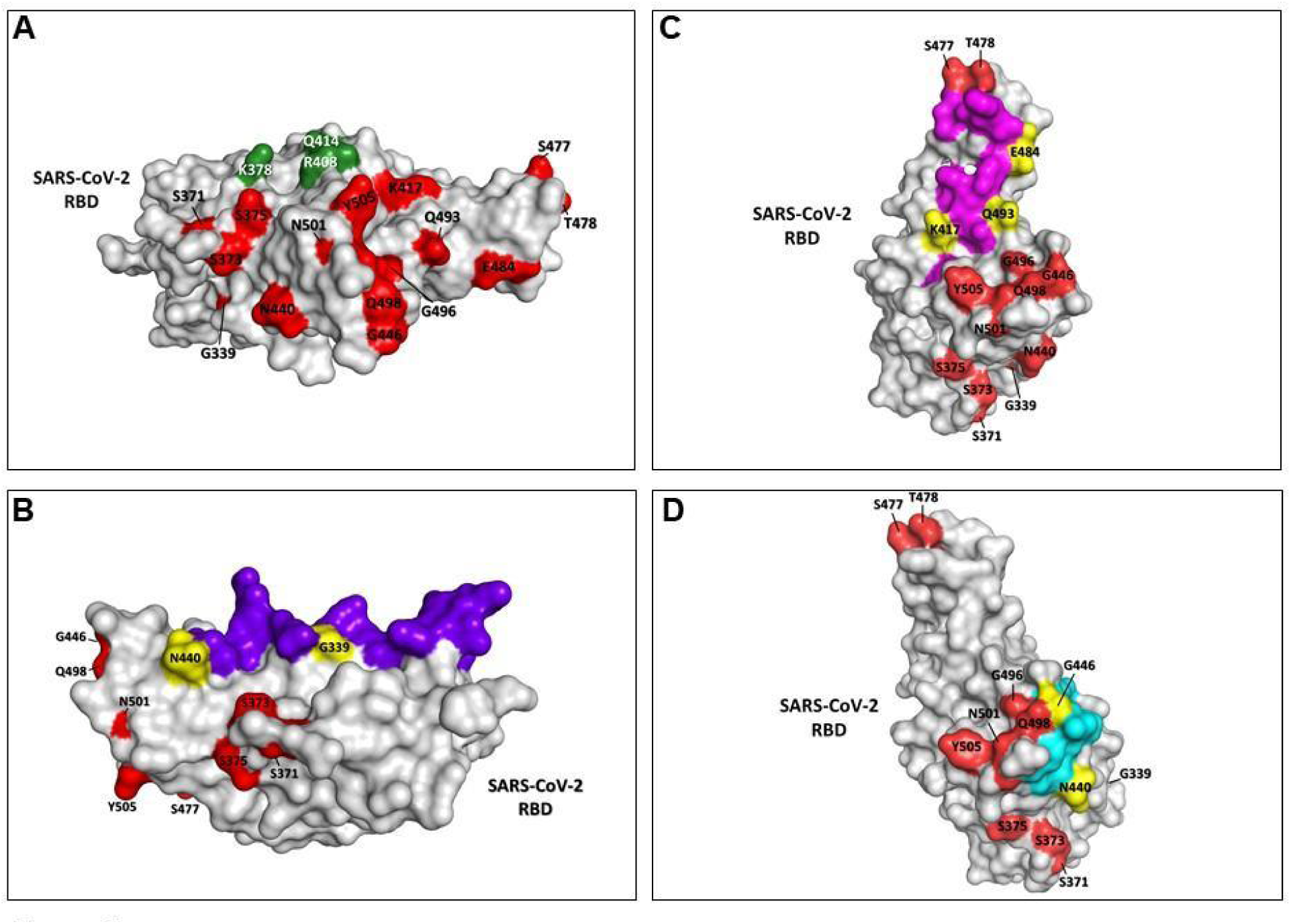
No critical binding residues for ABP-310 are mutated in the SARS-CoV-2 Omicron variant. **A-D**: Omicron variant mutation sites within the RBD are colored red, and residues that are both mutated in Omicron and within a given antibody’s epitope are colored yellow. **A**. ABP-310 critical binding residues of the SARS-CoV-2 spike RBD are colored dark green. **B**. S309 epitope residues are colored purple. The two S309 epitope residues mutated in Omicron (G339 and N440) are not critical binding residues. **C**. Casirivimab epitope residues are colored purple. The three casirivimab epitope residues mutated in Omicron (K417, E484, Q493) are critical binding residues. **D**. Imdevimab epitope residues are colored blue. The two imdevimab epitope residues mutated in Omicron (N440 and G446) are critical binding residues.

There appear to be no critical binding residues for S309 affected by Omicron mutations (Starr, 2021b) (**Fig. 5, Fig. 6**), as reflected by its retention of neutralizing activity against this variant (**Fig. 3B**) (GSK, 2021; Camaroni, 2021).

## Discussion

Here we report the identification of ABP-310, a neutralizing antibody binding a conserved epitope with potent activity against all current SARS-CoV-2 variants of concern, including Omicron, as well as neutralizing activity against SARS-CoV, demonstrating the ability to bind a conserved epitope and neutralize a related coronavirus (**Figs. 3 and 4, Extended Table 1, Extended Table 2**). Similarly, sotrovimab neutralized all SARS-CoV-2 variants and SARS-CoV, albeit at generally decreased potencies compared to ABP-310 (**Figs. 3B and 4, Extended Table 1, Extended Table 2**). Conversely, casirivimab and imdevimab were not able to neutralize all SARS-CoV-2 variants tested, failing against Omicron (**Fig. 3A**), and, in contrast to ABP-310 and sotrovimab, displayed little or no activity against SARS-CoV (**Fig. 4, Extended Table 1, Extended Table 2**). The failure of casirivimab, imdevimab (**Fig. 3A, Extended Table 1, Extended Table 2**), bamlanivimab, and estevimab to neutralize the Omicron variant (Cao, 2021; Planas, 2021) leaves only one effective SARS-CoV-2 nAb currently under EUA, highlighting the need for the development of SARS-CoV-2 nAb-based therapeutics more resistant to mutant escape.

The nAb described herein has several advantages over many existing SARS-CoV-2 nAb therapeutics. Unlike casirivimab and imdevimab, ABP-310 neutralizes all current and past SARS-CoV-2 variants tested, including Omicron, and also binds a conserved epitope, as shown by multiple sequence alignment with related coronaviruses (**Fig. 5**) and functionally demonstrated by the neutralization of the related coronavirus SARS-CoV (**Fig. 4**). Both ABP-310 and sotrovimab were to neutralize all of the SARS-CoV-2 variants tested as well as SARS-CoV, although ABP-310 did so with a mean 10.3-fold greater potency for SARS-CoV-2 variants and 2.1-fold greater potency for SARS-CoV than sotrovimab (**Fig. 3 and 4, Extended Table 1, Extended Table 2**). Our findings that of all the nAbs tested, only the conserved epitope binding antibodies ABP-310 and sotrovimab were able to neutralize the Omicron variant (**Fig. 3, Extended Table 1, Extended Table 2**), are consistent with previous results showing that nAbs binding conserved epitopes are less susceptible to Omicron escape (Cao, 2021). The conserved nature of the ABP-310 epitope also suggests that this antibody may be effective against other betacoronaviruses, including those yet to evolve, potentially contributing to an early therapeutic intervention during a future pandemic. The weak but detectable neutralizing activity observed for casirivimab against SARS-CoV (**Fig. 4**) is curious in light of the lack of observed binding of casirivimab to the SARS-CoV spike protein (**Fig. 1**). It should be pointed out that the maximum concentrations examined were 5 µg/ml and 100 µg/ml for the Luminex binding and pseudovirus neutralization assays, respectively, thus the lack of binding observed in the Luminex assay may be a result of testing insufficiently high concentrations of antibody. However, as the SARS-CoV neutralizing IC_50_ for casirivimab is at least 11-fold less potent than for soluble ACE2 (data not shown), it is questionable if casirivimab would be an effective therapeutic against SARS-CoV or other related coronaviruses existing or yet to evolve. In contrast, the potent ability of ABP-310 to neutralize SARS-CoV is reflected in its 49-fold more potent neutralizing IC_50_ than soluble ACE2 (data not shown).

It has been suggested that the widespread use of transgenic cell lines overexpressing ACE2 in viral neutralization assays can underestimate the potency and efficacy of antibodies that bind the SARS-CoV-2 spike protein RBD outside the receptor binding motif (RBM) such as sotrovimab, especially in light of the claims that ACE2-expression is low in the lower respiratory tract. Rather, the use of cells with low ACE2 expression, such as the African green monkey kidney cell-derived VeroE6 cell line has been suggested (Lempp, 2021). While it is true that ACE2-expression is restricted to relatively few cell types in the lower respiratory tract, most notably alveolar type II (AT2) cells (**Hamming, 2004**) as well as alveolar type I cells, airway epithelial cells, fibroblasts, endothelial cells, and macrophages (Zhao, 2020), AT2 cells express high levels of ACE2 and Type I and II Interferons (Ziegler, 2020) and IL-1β (Chen, 2019) can induce ACE2 expression in a wide array of respiratory tissue cells. In addition, there is a general gradient of ACE2 cellular expression running from higher in the upper respiratory tract to lower in the lower respiratory tract (Hou, 2020). It is possible that the upper respiratory tract may be an initial site of infection, with aspiration or other mechanisms bringing virus into the lower respiratory tract, infecting AT2 cells and other cell types whose ACE2 expression can be upregulated through inflammatory mediators such as Interferons and IL-1β. Indeed, the ability of SARS-CoV-2 to be able to infect cells of the upper respiratory system has been demonstrated for the Omicron variant (Diamond, 2021; Peacock, 2022). Collectively, these findings demonstrate that there are plentiful high ACE2-expressing cells as targets for viral infection in both the upper and lower respiratory tracts. In addition to the low expression levels of ACE2 on VeroE6 cells, African green monkey ACE2 is only 89% identical to human ACE2 (Sheahan, 2008) suggesting there may be differences in the efficiency of human-tropic SARS-CoV-2 to infect these cells. This brings into question the suitability of using ACE2 low expressing cells, especially ACE2 low expressing cells of other species such as VeroE6 cells. An examination of the effect of ACE2 expression levels on cells used in *in vitro* viral neutralization assays and found that while there was an inverse correlation between S309 neutralization and ACE2 expression on cells used in a VSV-based, SARS-CoV-2 spike protein pseudovirus neutralization assay, there was no such correlation when using a live virus assay, with the exception of HEK-293T cells transgenic for ACE2 (Lempp, 2021). In a separate study, there was no difference found in S309 neutralization efficacy (percent maximum neutralization) or potency (IC_50_s) in SARS-CoV-2 live virus neutralization assays using parental VeroE6 cells or transgenic human ACE2-VeroE6 cells (Chen, 2021). Similarly, in our lentiviral-based SARS-CoV-2 spike protein pseudovirus neutralization assay using ACE2-transgenic HEK-293T cells, sotrovimab achieved a mean maximum neutralization of 92% across the SARS-CoV-2 variants tested with activity against most variants being greater than 98% (**Fig. 3**), consistent with published live virus assays and pseudovirus assays performed with VeroE6 cells where maximum neutralization also approached 100% (Chen, 2021; Lempp, 2021). As the use of ACE2-transgenic cell lines in live virus neutralization assays (Chen, 2021; Lempp, 2021; Planas, 2021) or our lentiviral-based pseudovirus neutralization assay does not appear to adversely affect the efficacy or potency of S309 or sotrovimab, the choice of pseudovirus system (e.g., VSV, MLV, Lentiviral) for neutralization assays appears to be crucial for effective *in vitro* evaluation of nAbs, especially those binding epitopes outside the RBM. Taken together, the presence of high ACE2-expressing cells throughout the upper and lower respiratory tracts as well as the ability of sotrovimab to potently neutralize virus using target cells expressing high levels of ACE2 suggests that the use of target cell lines with low ACE2 expression, instead of underestimating, may be overestimating the potency and efficiency of non-RBM binding antibodies. In practice, so long as the overestimation is modest, the effect on clinical efficacy may be negligible. For sotromivab, the 500mg dose (FDA Fact sheet for sotrovimab) would be expected to result in peak plasma concentrations in a 60kg person of approximately 119 µg/ml after intravenous dosing, which is 40-250-fold higher than reported IC_50_ values against wild type SARS-CoV-2 from a wide variety of assay formats using different pseudoviral and authentic live viral systems (Pinto, 2020; Chen, 2021) (**Extended Table 1**).

In summary, we describe a SARS-CoV-2 neutralizing antibody, ABP-310, with broad SARS-CoV-2 variant coverage (including Omicron), conserved epitope binding, and ability to neutralize SARS-CoV. We believe that the broad neutralizing ability within SARS-CoV-2 variants shows the potential of ABP-310 as an effective therapeutic for treatment or prophylaxis of SARS-CoV-2 infection. Additionally, given its conserved epitope-binding and cross-species viral neutralization, it may also serve as a frontline therapeutic against yet to evolve lineage B betacoronaviruses.

## Methods and Materials

### Comparator control antibody production

Casirivimab and imdevimab (ProteoGenix, Schiltigheim, France) as well as sotrovimab (Abpro, Woburn, MA, USA) were produced at the aforementioned institutions using the sequence of each antibody as obtained from The United States Adopted Name Council of the American Medical Association.

### ACE2 competition assay

The competition reaction was tested by using the SARS-CoV-2 variant inhibitor screening kit (R&D VANC00) according to the manufacturer’s instruction to assess whether the selected antibodies can compete with ACE2 for RBD binding. Briefly, 96 well plates were coated with His-Tag capture antibody and incubated at 4°C overnight then blocked with 1% BSA in PBS for 1 hour at 37°C. Afterward recombinant SARS-Cov-2 wild type (Wuhan) spike protein RBD or mutant variants were immobilized onto His-Tag capture antibody-coated plates for an additional of 1 hour at room temperature. Serial dilutions of the selected antibodies were loaded in duplicates and incubated for 1 hour at room temperature, followed by the addition of biotinylated human ACE2. Then Streptavidin-HRP was added, and the plates were developed using substrate solution, followed by sulfuric acid addition to stop the reaction. The plates were read at 450 nm by a SpectraMax iD3 Multi-Mode Microplate Reader manufactured by Molecular Devices to determine the results.

To calculate the percentage of ACE2 binding, the average nonspecific binding was subtracted from each point and divided by RBD only control value and multiplied by 100.

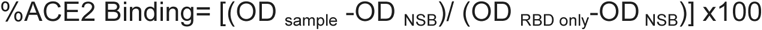

Results were analyzed in GraphPad PRISM Version 9.3.0.

### Bead-based multiplex assay method for SARS-CoV-2 spike protein binding

A custom assay was designed using Magplex-Avidin microspheres (Luminex), which were coated in anti-HIS antibody [Biotin] (GenScript A00613, mouse IgG1k clone 6G2A9) at 2.5 μg/mL, washed, and then each microsphere was coated with 2.5 μg/mL of either SARS-CoV-1 S1-HIS, MERS S1-HIS, and SARS-CoV-2 S1-HIS or SARS-CoV-2 RBD-HIS variant proteins (Sino Biological) to create bead stocks. For the antibody binding assay, test antibodies were diluted in PBS/1% BSA starting at 10 μg/mL (2x concentrations) and then five-fold on U-bottom plates, for a total of eight dilutions. 50 μL/well of antibody was added to a 96-well plate (Greiner Bio-One 655096). A master stock of microspheres was created in PBS/1% BSA to allow 2000 beads/analyte/well, vortexed, and 50 μL/well was added to the assay plate. Antibodies were incubated with microspheres for 30 minutes at RT with shaking at 300 rpm, washed 3x with PBS, then incubated with Goat anti-Human IgG Fc Secondary Antibody PE, eBioscience (minimal cross-reactivity to bovine/horse/mouse serum proteins)(Thermo Fisher Scientific 12-4998-82) diluted 1:200 in PBS/1% BSAfor 30 minutes.). Wells were washed 3x with PBS, and the plate was read on the Magpix (Luminex) instrument. Median Fluorescence Intensity (MFI) was used to quantify binding of antibodies to each protein in the panel. See **Extended Table 3** for the HIS-tagged proteins used in the assay. Results were analyzed in GraphPad PRISM Version 9.3.0.

### Pseudovirus neutralization assays

SARS-CoV-2 antibodies were tested for neutralization of pseudovirus particles expressing the wild type spike protein of SARS-CoV, SARS-CoV-2, or variants thereof (Integral Molecular, see **Extended Table 4**) according to manufacturer’s protocol. Briefly, the recommended amount of particles/well was incubated in DMEM + 10% FBS with varying amounts of serially-diluted antibody for 1 hour at 37°C, and then 20,000 ACE2-HEK293T cells (Integral Molecular, see **Extended Table 5**) were added. Neutralization of infection was assessed using Renilla-Glo Luciferase Assay kit (Promega) after an incubation of 48-72 hours. Data was collected by a SpectraMax iD3 Multi-Mode Microplate Reader manufactured by Molecular Devices. Results were analyzed in GraphPad PRISM Version 9.3.0 and IC_50_ values were determined for individual antibodies.

### Structural analysis

The figures were rendered using PyMOL Version 2.4.1 (The PyMOL Molecular GraphicsSystem; http://www.pymol.Org). The SARS-CoV-2 spike RBDs were presented in surface form. The model of critical binding residues for ABP-310 on the SARS-CoV-2 spike RBD was generated using the RBD of PDB ID 7JX3 (wwpdb.org) as a template. The structural models of the SARS-CoV-2 spike RBD binding footprints for REGN10933 (casirivimab) and REGN10987(imdevimab) were obtained using PDB ID 6XDG, and for S309 using PDB ID 7JX3 (wwpdb.org).

### Multiple alignment of related lineage B betacoronavirus spike protein sequences

Multiple alignment was made using Clustal Omega (Madeira, 2019) and image rendered using Geneious software (Kearse, 2012). Clades classification and sequences in the multiple alignment analysis were previously published (Letko, 2020).

### Alanine Scanning via shotgun mutagenesis

Shotgun Mutagenesis epitope mapping services were provided by Integral Molecular (Philadelphia, PA) as previously described (Davidson, 2014). Briefly, a mutation library of the target protein was created by high-throughput, site-directed mutagenesis. Each residue was individually mutated to alanine, with alanine codons mutated to serine. The mutant library was arrayed in 384-well microplates and transiently transfected into HEK-293T cells. Following transfection, cells were incubated with the indicated antibodies at concentrations pre-determined using an independent immunofluorescence titration curve on wild type protein. MAbs were detected using an Alexa Fluor 488-conjugated secondary antibody and mean cellular fluorescence was determined using Intellicyt iQue flow cytometry platform. Mutated residues were identified as being critical to the MAb epitope if they did not support the reactivity of the test MAb but did support the reactivity of the reference MAb. This counterscreen strategy facilitates the exclusion of mutants that are locally misfolded or that have an expression defect.

## Author Contributions

Conceptualization: A.J.P., S.P.M., S.E., J.L., S.W.; Methodology: A.J.P., S.E., J.L., and S.P.M.; Investigation: A.J.P., S.E., L.Z., and O.B.; Data Analysis: A.J.P., S.P.M., S.E., J.L., L.D., C.Z., L.Z., and O.B.; Writing-original draft: S.P.M.; Writing-review and editing: S.P.M, A.J.P., S.E., J.L., E.C., L.D., C.Z., L.Z., O.B., G.A., and AK.; Supervision: S.P.M., A.J.P., G.A., J.L., and E.C.

## Figure and Table Legends

**Extended Table 1.**
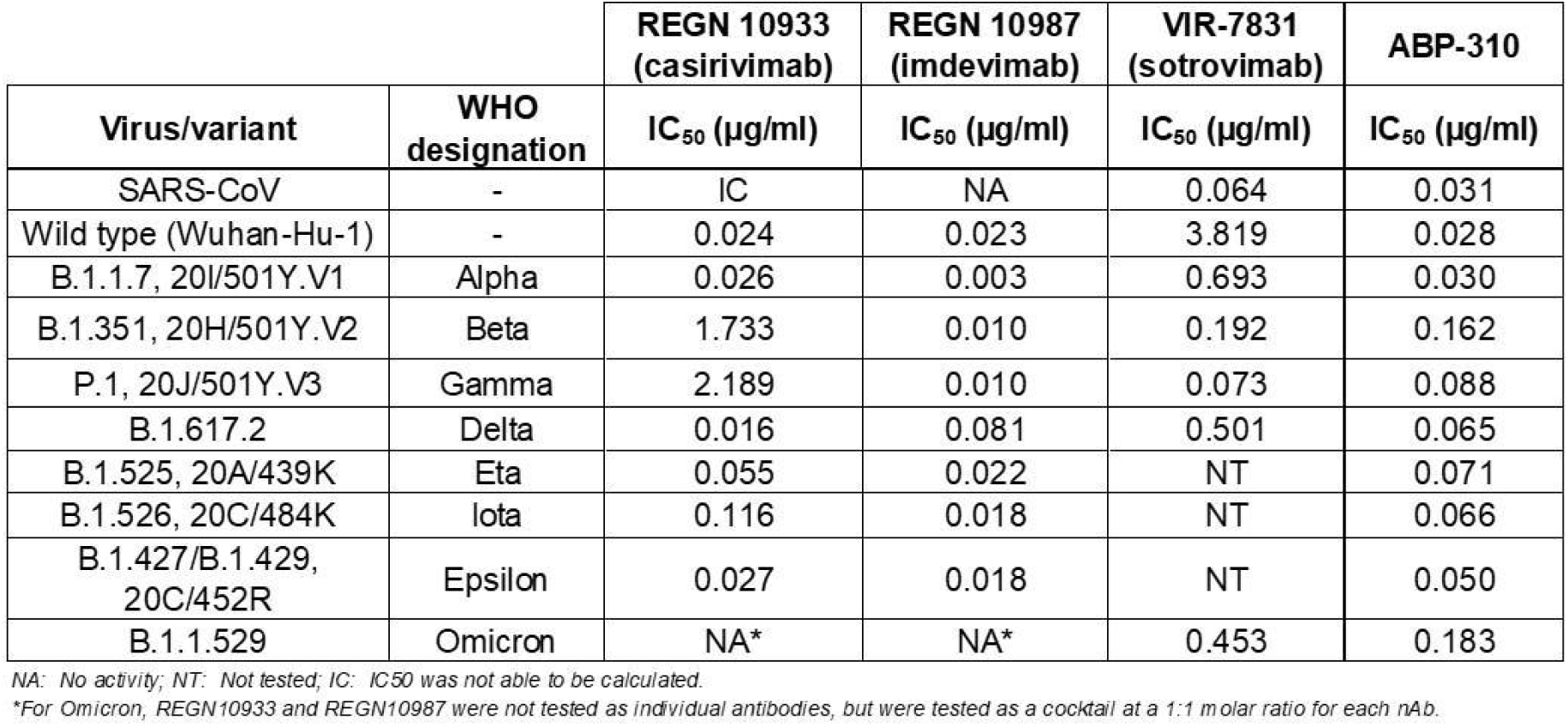
Neutralization IC50s of neutralizing antibodies against indicated pseudotyped viruses. Neutralizing IC_50_s for antibodies against SARS-CoV and indicated SARS-CoV-2 variants.

**Extended Table 2.** Neutralization IC_90_s of neutralizing antibodies against indicated pseudotyped viruses. Neutralizing IC_90_s for antibodies against SARS-CoV and indicated SARS-CoV-2 variants.

| Virus/variant | WHO designation | REGN 10933 (casirivimab) | REGN 10987 (imdevimab) | VIR-7831 (sotrovimab) | ABP-310 |
| --- | --- | --- | --- | --- | --- |
|  |  | IC<sub>90</sub> (µg/ml) | IC<sub>90</sub> (µg/ml) | IC<sub>90</sub> (µg/ml) | IC<sub>90</sub> (µg/ml) |
| SARS-CoV | - | IC | NA | <sup>†</sup> | 0.233 |
| Wild type | - | 0.136 | 0.319 | 138.900 | 0.163 |
| B.1.1.7, 20I/501Y.V1 | Alpha | 0.237 | 0.029 | 12.970 | 0.053 |
| B.1.351, 20H/501Y.V2 | Beta | 13.940 | 0.093 | 0.650 | 0.696 |
| P.1, 20J/501Y.V3 | Gamma | 27.330 | 0.060 | 0.515 | 0.551 |
| B.1.617.2 | Delta | 0.098 | 0.504 | 2.920 | 0.179 |
| B.1.525, 20A/439K | Eta | 0.480 | 0.275 | NT | 0.201 |
| B.1.526, 20C/484K | Iota | 1.217 | 0.217 | NT | 0.237 |
| B.1.427/B.1.429, 20C/452R | Epsilon | 0.189 | 0.228 | NT | 0.176 |
| B.1.1.529 | Omicron | NA* | NA* | 4.915 | 0.798 |

NA: No activity; NT: Not tested; <sup>†</sup> Sotrovimab did not achieve 90% neutralization for SARS-CoV. IC: IC<sub>50</sub> was not able to be calculated.
\*For Omicron, REGN10933 and REGN10987 were not tested as individual antibodies, but were tested as a cocktail at a 1:1 molar ratio for each nAb.

**Extended Table 3.**
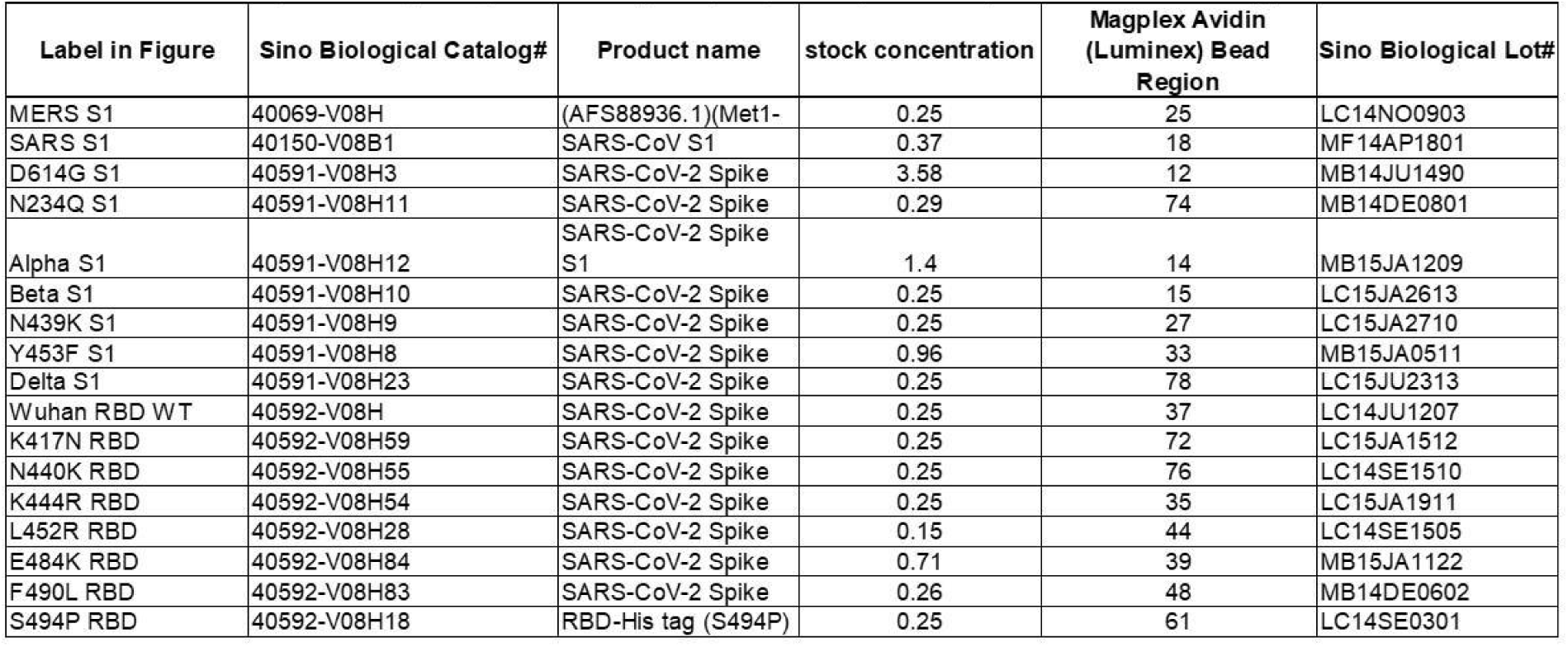
Spike proteins used in Luminex binding assays. Multiplex binding assay: Sino Biological spike protein (S1 or RBD, with C-terminal HIS-tag)

| Label in Figure | Sino Biological Catalog# | Product name | stock concentration | Magplex Avidin (Luminex) Bead Region | Sino Biological Lot# |
| --- | --- | --- | --- | --- | --- |
| MERS S1 | 40069-V08H | (AFS88936.1)(Met1- | 0.25 | 25 | LC14NO0903 |
| SARS S1 | 40150-V08B1 | SARS-CoV S1 | 0.37 | 18 | MF14AP1801 |
| D614G S1 | 40591-V08H3 | SARS-CoV-2 Spike | 3.58 | 12 | MB14JU1490 |
| N234Q S1 | 40591-V08H11 | SARS-CoV-2 Spike | 0.29 | 74 | MB14DE0801 |
| Alpha S1 | 40591-V08H12 | SARS-CoV-2 Spike S1 | 1.4 | 14 | MB15JA1209 |
| Beta S1 | 40591-V08H10 | SARS-CoV-2 Spike | 0.25 | 15 | LC15JA2613 |
| N439K S1 | 40591-V08H9 | SARS-CoV-2 Spike | 0.25 | 27 | LC15JA2710 |
| Y453F S1 | 40591-V08H8 | SARS-CoV-2 Spike | 0.96 | 33 | MB15JA0511 |
| Delta S1 | 40591-V08H23 | SARS-CoV-2 Spike | 0.25 | 78 | LC15JU2313 |
| Wuhan RBD WT | 40592-V08H | SARS-CoV-2 Spike | 0.25 | 37 | LC14JU1207 |
| K417N RBD | 40592-V08H59 | SARS-CoV-2 Spike | 0.25 | 72 | LC15JA1512 |
| N440K RBD | 40592-V08H55 | SARS-CoV-2 Spike | 0.25 | 76 | LC14SE1510 |
| K444R RBD | 40592-V08H54 | SARS-CoV-2 Spike | 0.25 | 35 | LC15JA1911 |
| L452R RBD | 40592-V08H28 | SARS-CoV-2 Spike | 0.15 | 44 | LC14SE1505 |
| E484K RBD | 40592-V08H84 | SARS-CoV-2 Spike | 0.71 | 39 | MB15JA1122 |
| F490L RBD | 40592-V08H83 | SARS-CoV-2 Spike | 0.26 | 48 | MB14DE0602 |
| S494P RBD | 40592-V08H18 | RBD-His tag (S494P) | 0.25 | 61 | LC14SE0301 |

**Extended Table 4.** Pseudoviral particles for neutralization assays expressing coronavirus spike protein.

| Variant pseudoviral particle | Product (Integral Molecular) | Strain/variant name (Integral Molecular) | WHO | Mutations | D614G? |
| --- | --- | --- | --- | --- | --- |
| SARS-CoV-1 | RVP-801L | SARS-CoV-1 Urbani | - | - | No |
| SARS-CoV-2 ("WT") | RVP-701L | Wuhan-Hu-1 | - | Reference seq (GenBank QHD43416.1) | No |
| D614G | RVP-702L | D614G B.1, 20A | - | D614G | Yes |
| Eta (Nigerian) | RVP-723L | Nigerian/European variant B.1.525, 20A/439K | Eta | Q52R, ΔH69/V70, ΔY144, E484K, Q677H, F888L | Yes |
| Iota (New York) | RVP-726L | New York variant B.1.526, 20C/484K | Iota | L5F, T95I, D253G, E484K, A701V | Yes |
| Beta (South African) | RVP-724L | South African variant Δ3 B.1.351, 20H/501Y.V2 | Beta | L18F, D80A, D215G, ΔL242/A243/L244, R246I, K417N, N501Y, E484K, A701V | Yes |
| Indian "double mutant" | RVP-727L | Indian variant B.1.617, RBD mutations only ("double mutant") | - | L452R, E484Q | Yes |
| Epsilon (California) | RVP-713L | Californian variant B.1.427/B.1.429, 20C/452R | Epsilon | S13I, W152C, L452R | Yes |
| Alpha (U.K.) | RVP-706L | UK variant B.1.1.7, 20I/501Y.V1 | Alpha | ΔH69/V70, ΔY144, N501Y, A570D, P681H, T716I, S982A, D1118H | Yes |
| Gamma (Brazil/Japan) | RVP-708L | Brazilian variant P.1, 20J/501Y.V3 | Gamma | L18F, T20N, P26S, D138Y, R190S, K417T, E484K, N501Y, H655Y, T1027I, V1176F | Yes |
| Delta | RVP-763L | Indian variant B.1.617.2 | Delta | T19R, G142D, del156/157, R158G, L452R, T478K, P681R, D950N | Yes |
| Omicron | RVP-768L | Omicron Variant B.1.1.529 | Omicron | A67V, Δ69-70, T95I, G142D/Δ143-145, Δ211/L212I, ins214EPE, G339D, S371L, S373P, S375F, K417N, N440K, G446S, S477N, T478K, E484A, Q493R, G496S, Q498R, N501Y, Y505H, T547K, D614G, H655Y, N679K, P681H, N764K, D796Y, N856K, Q954H, N969K, L981F | Yes |

**Extended Table 5.**
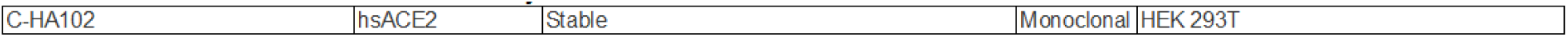
Cell line used for pseudovirus assays.

|  |  |  |  |  |
| --- | --- | --- | --- | --- |
| C-HA102 | hsACE2 | Stable | Monoclonal | HEK 293T |

